# Dendritic plateau potentials can process spike sequences across multiple time-scales

**DOI:** 10.1101/690792

**Authors:** Johannes Leugering, Pascal Nieters, Gordon Pipa

## Abstract

The brain constantly processes information encoded in temporal sequences of spiking activity. This sequential activity emerges from sensory inputs as well as from the brain’s own recurrent connectivity and spans multiple dynamically changing timescales. Decoding the temporal order of spiking activity across these varying timescales is a critical function of the brain, but we do not yet understand its neural implementation. The problem is, that the passive dynamics of neural membrane potentials occur on a short millisecond timescale, whereas many cognitive tasks require the integration of information across much slower behavioral timescales. However, actively generated dendritic plateau potentials do occur on such longer timescales, and their essential role for many aspects of cognition has been firmly established by recent experiments. Here, we build on these discoveries and propose a new model of neural computation that emerges from the interaction of localized plateau potentials across a functionally compartmentalized dendritic tree. We show how this interaction offers a robust solution to the timing invariant detection and processing of sequential spike patterns in single neurons. Stochastic synaptic transmission complements the deterministic all-or-none plateau process and improves information transmission by allowing ensembles of neurons to produce graded responses to continuous combinations of features. We found that networks of such neurons can solve highly complex sequence detection tasks by breaking down long inputs into sequences of shorter, random features that can be classified reliably. These results suggest that active dendritic processes are fundamental to neural computation.

## Introduction

The ability to recognize patterns of spiking activity across multiple timescales is a remarkable and crucial function of the brain. For example, consider a rodent searching for food. The receptive fields of its place and grid cells tile a spatial map of the environment and encode its current position by their respective population activities [1, 2]. To navigate successfully, the animal needs to know not only its present location, but also the path it took to get there. This path is encoded in the sequential firing of place and grid cells on behavioral timescales that can span hundreds of milliseconds or more [3]. While the exact timing of these spikes depends on the speed of the animal, the chosen path through a space is encoded in the sequential order in which these cells are activated. As another perhaps more familiar modality, consider natural language: We can understand the same utterances over a wide variety of speech tempos and rhythms, provided that the order of specific sounds is preserved. In auditory cortex, for example, sequences of brief phonemes encode longer syllables which can last for around 200 ms [4]. Early olfactory areas represent odors as ordered sequences on a similar timescale [5, 6]. And visual cortex can learn to reproduce sequential activation patterns on cue, but on a compressed timescale [7, 8]. The sum of these and other findings suggest that sequential patterns of spiking activity, the precise timing of which can vary across timescales of up to several hundred milliseconds, are a ubiquitous occurrence in the brain. Therefore, the ability to detect and decode such sequences of neural activity is likely crucial for complex behavior in humans and animals alike [9]. But how neural circuits implement this is currently unknown.

Point-neurons such as leaky integrate-and-fire models operate on the fixed timescale of passive membrane potential dynamics, which is typically on the order of tens of milliseconds or less [10]. Since this is at least one order of magnitude shorter than the behavioral timescales we are interested in here, there must be additional working memory to bridge the silent periods between successive bouts of neural activity in longer sequences. Often, this longer memory is attributed to network-level effects such as slow recurrent dynamics [11] or fast synaptic plasticity [12]. However, computational properties of neural systems are also dependent on the structure and dynamics in the neural dendrites of single neurons [13], which make up the vast majority of neural tissue [14]. We are particularly interested in dendritic plateau potentials [15], a localized all-or-none response in the membrane potential of pyramidal neurons’ dendrites. Plateaus can be triggered by coherent volleys of spikes, and the resulting elevated membrane potentials are maintained for up to hundreds of milliseconds [16]. These actively generated events are ubiquitous throughout the brain [17, 18], and recent in-vivo experiments link the prolonged high internal Ca^2+^ concentrations caused by dendritic plateaus to various cognitive tasks [19, 20, 21, 22]. As previously suggested [23], this implies that dendritic plateau potentials may be the key missing ingredient in current models of sequence processing in the brain, but their precise role is still debated.

Our main idea is that dendritic plateaus provide the long-lasting memory traces needed for sequence processing: By generating a plateau potential, the dendrite quickly reacts to the detection of a stimulus, i.e. coherent synaptic input, and retains a long-lasting memory trace of this event in the local membrane potential. The key mechanism for detecting sequences of such stimuli on behavioral timescales is the interaction of plateau potentials across nearby segments, i.e. the depolarizing effect they exert on neighboring segments. We propose that localized dendritic plateaus thus turn the complex dendritic trees of single neurons into reliable sequence processors with long temporospatial, yet timing-invariant receptive fields.

In the following, we first derive from recent biological observations a conceptual model of dendritic computation that is built on this interaction of plateau potentials. Then we demonstrate how this mode of computation allows single neurons and networks to recognize complex sequences across various timescales and modalities.

### Stochastic generation and inhibition of localized dendritic plateau potentials

Most of a cortical pyramidal neuron’s excitatory synaptic inputs terminate on dendritic spines [24], where they activate both AMPAr and NMDAr gated ion channels via glutamate-carrying vesicles. The AMPAr channel opens immediately, which leads to a brief depolarization in the corresponding spine referred to as the *excitatory post-synaptic potential* (EPSP) [25]. The NMDAr channel, however, is blocked by an Mg^+^ ion that must first be displaced by sufficient depolarization of the postsynaptic membrane to become conductive [26, 27]. This requires the cumulative effect of coincident EPSPs in multiple nearby spines [28]. Experimental as well as simulation studies report that a volley of 4-20 or even up to 50 spikes within a window of 1ms to 4ms is needed to provide sufficient depolarization, depending on the location along the dendritic tree [28, 29, 30, 31]. The opening of NMDAr channels triggers a massive influx of different ionic currents that lead to a full depolarization of the local dendritic membrane potential. Although the isolated NMDAr response itself is reported to only last on the order of around 25 ms [32], in vivo recordings reveal that voltage-gated channels in the dendritic membrane [33] prolong this effect, resulting in an actively maintained depolarization that can last from tens to hundreds of milliseconds [34]. This prolonged depolarization is called a dendritic plateau potential. Two further aspects of the interaction between dendritic plateaus and synapses are noteworthy. Firstly, spikes in presynaptic neurons only lead to the required glutamate release some of the time with a synapse specific probability [35, 36]. In hippocampal synapses for example, the median release probability is only 0.22 [37]. Secondly, the activation of inhibitory GABA_*A*_ and GABA_*B*_ synapses can strongly interfere with the dendritic plateau process and outright stop or prevent plateau generation [38, 39, 40]. The importance of inhibitory synapses is further emphasized by their locations, which are often in critical positions to control dendritic excitability [41] and gate specific dendritic activity from reaching the soma [42].

### Functional compartmentalization of dendritic trees

Plateau potentials remain localized to specific sites in the dendritic tree because their generation and maintenance requires the binding of external glutamate to NMDAr channels [43]. Therefore, the structure of dendritic arbors, which has long been conjectured to play an important role for neural computation, is crucial for plateau computation. For example, functional subunits emerge in the dendrites of various types of retinal ganglion cells due to impedance mismatches at branching points in the dendritic tree [44]. These are regions of roughly equal local membrane potentials throughout, but they are only weakly coupled to neighboring regions. Other experiments in rats confirmed that thin dendrites in neocortical pyramidal neurons can act as independent computational subunits and provide neurons with additional non-linear integration sites [45]. This behavior is not limited to pyramidal neurons, but rather appears to be a general principle that can be found in various forms across different cell types. For example, Purkinje cells in the cerebellum also generate localized Ca^2+^ events in response to coincident input on individual dendrite segments [46, 47], and thalamo-cortical neurons respond to strong synaptic input with localized plateaus in distal dendritic branches [48]. In some neurons, functional subunits can be identified with individual dendritic branches [49]. More generally, these subunits can also stretch across multiple nearby branches if they are sufficiently strongly coupled so that coherent synaptic input across the branches can trigger local, regenerative events such as plateau potentials [50]. We view dendrites as complex structures composed of functional subunits in the latter sense and will refer to them as dendrite segments.^2^

Neural cable theory predicts an asymmetric passive propagation of membrane potentials throughout the dendrite [51, 52]. In the anterograde direction, the signal attenuation is generally so strong that synaptic input onto thin apical dendrites has little measurable effect of the membrane potential at the soma [53, 33]. On the one hand, this may suggest that superlinear NMDAr responses along the dendrite serve to boost distal input signals [54, 55]. On the other hand, plateau potentials are also subject to attenuation along the dendritic cable and thus only have a moderately depolarizing effect on their immediate neighborhood [56]. This can effectively raise the local resting potential of a neighboring segment closer to the soma and thus lower the amount of coinciding spikes required to initiate a plateau potential in it [16].

### From dendritic nonlinearity to dendritic computation

Neither stochastically generated active dendritic processes nor the structure of the dendritic tree are considered in most computational models of spiking neurons, although there is lively debate about the appropriate level of abstraction [57]. Some authors argue that the dendritic integration of spikes can be largely explained by a linear stochastic model with one additional non-linear term [58]. Others contend that the computational function of the dendritic tree is best captured by a non-linear hidden layer in a neural network model [59], whereas modeling the temporal dynamics of the membrane potential would require significantly more complex temporally convolutional deep neural networks [60]. But the long-lasting all-or-none response characteristic of dendritic plateaus is not captured by either of these model classes. We therefore take a different approach and focus entirely on dendritic plateaus and their interactions in the dendrite. Since only nearby synapses cooperate to trigger plateau potentials, correlated spikes have to arrive at the same dendrite segment at the same time to effectively drive plateau generation [30]. This has been confirmed in experiments [61], suggesting that some information in the brain is conveyed by highly synchronized spike volleys that target individual dendrite segments [62] and trigger dendritic plateaus there. We argue that the interaction of these plateaus gives rise to dendritic computation on behavioral timescales. This approach avoids the complexities of non-linear membrane dynamics altogether, and it offers a clearer picture of the underlying computational mechanism that we believe may be vital to sequence processing, in particular.

### A computational model for dendritic plateau computation

From the biological observations outlined above, we derive a simple, qualitative model of active dendrites. At the core of this model lies the interaction of two types of events on distinct timescales — short, spike-triggered EPSPs and long, actively generated dendritic plateau potentials — in a tree structure of dendrite segments. We define a *segment* as a minimal functional subunit of the dendritic tree, a single physical branch or multiple branches, that behave as one electrically isolated integration site. Synaptic inputs to a segment can therefore cooperatively generate a plateau potential that stays confined to the segment. These dendrite segments form a tree structure with the soma at its root and thin, distal branchlets as leaves.

First, consider the function of one individual dendrite segment *i* in more detail (see Fig. 1 **(C)** for a schematic). We distinguish excitatory and inhibitory synapses, which respectively produce excitatory (EPSPs) and inhibitory (IPSPs) postsynaptic potentials. An excitatory synapse from neuron *k* to segment *i* only successfully transmits each spike with probability *p*_*i,k*_. If the synapse transmits the spike, it induces an EPSP *κ*_*E*_(*t*) with duration *τ*_E_ and a magnitude *w*_*i,k*_, which depends on the synaptic efficacy. Likewise for an inhibitory synapse, only that the duration *τ*_I_ of the IPSP is typically slightly longer. We model the shape of the post-synaptic potentials by rectangular pulses:

**Figure 1:**
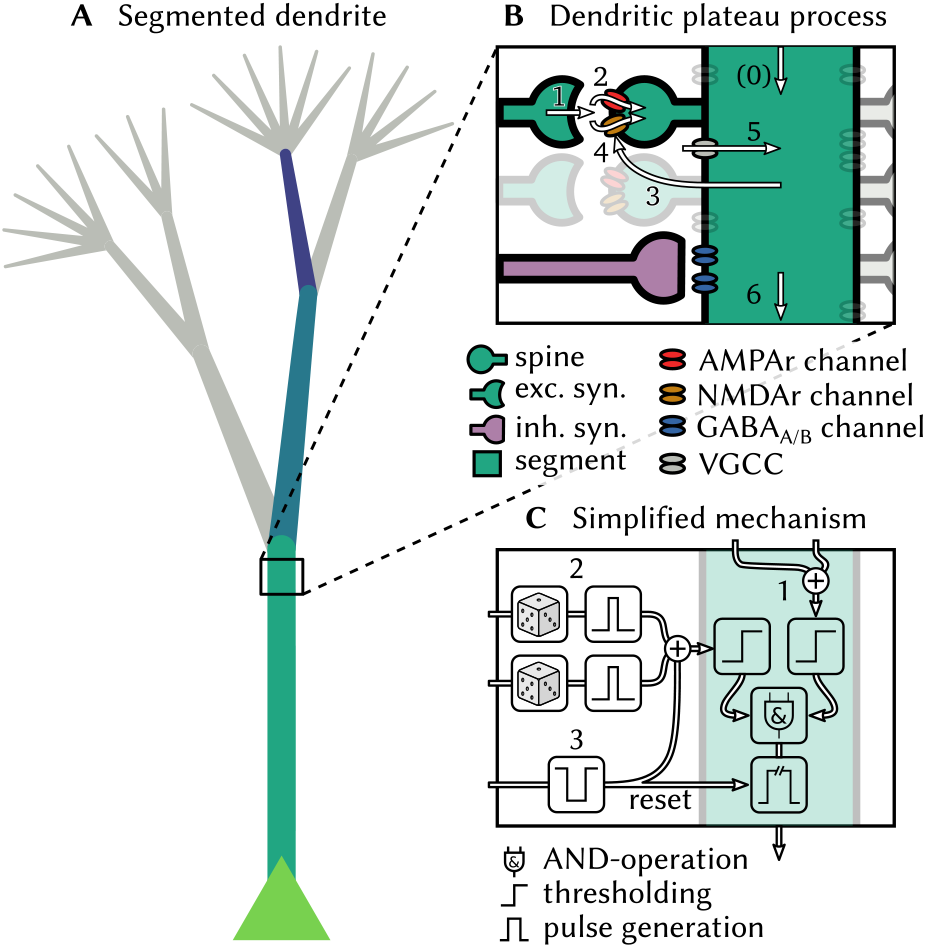
Generation and interaction of dendritic plateau potentials. **(A)** A stylized neuron with dendritic arbor. **(B)**Summary of the biological processes involved. A spike (1) releases glutamate, which opens AMPAr-gated ion-channels that depolarize the post-synaptic spine and cause an EPSP (2). If sufficiently many EPSPs coincide with up-stream dendritic input (0), the local membrane potentials rises (3) and NMDAr-gated ion-channels become de-inactivated, causing a further localized depolarization (4). Additional voltage-gated calcium channels can amplify and prolong this process (5) and cause a plateau potential, which can in turn moderately depolarize the parent segment (6 & 0). **(C)** An algorithmic approximation of the biological mechanisms. If a dendrite segment is depolarized by sufficiently strong input from its child segments (1) and receives sufficiently strong excitatory input from its stochastic synapses (2), a local plateau potential is initiated. If the plateau is not interrupted by shunting inhibitory input (3), it depolarlizes the parent segment for an extended period of time.

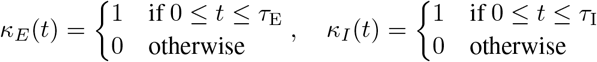

We use exc_*i*_ and inh_*i*_ to represent the set of excitatory and inhibitory neurons targeting segment *i*, we denote the time of the *m*^th^ spike by neuron *k* with 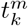, and introduce the i.i.d. random variables 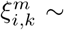 Bernoulli(*p*_*i,k*_) to simplify notation. We can then define the combined effect of excitatory as well as inhibitory input for segment *i*^3^:

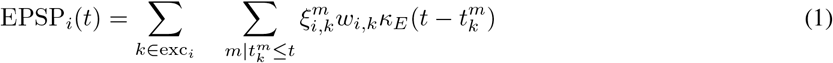

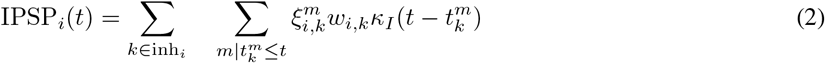

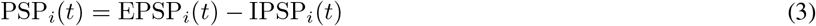

One of the necessary preconditions for generating a dendritic plateau potential is a sufficiently strong net depolarization of the dendrite by synaptic input. This means that the coincidence of multiple synchronous spikes caused depolarization larger than a segment-specific synaptic threshold TS_*i*_. In thin dendrite branchlets, which are the leaf nodes of our tree structure, this is sufficient to trigger a plateau potential. But in the general case, additional depolarizing input from dendritic child branches is required. Here, we are only interested in the large depolarizing effects that actively generated plateau potentials have on directly adjacent segments, and we ignore the much weaker passive propagation of sub-threshold voltages along the dendrite.

We therefore introduce additional notation: child_*i*_ denotes the set of the direct children of segment *i* (if any), and O_*k*_(*t*), *k* ∈ child_*i*_ is the effect that the child segment *k* exerts on *i* at time *t*. Just like we did for the post-synaptic potentials, we can then define the total *dendritic input* D_*i*_(*t*) into segment *i*:

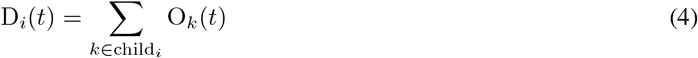

The segment-specific dendritic threshold TD_*i*_ determines how much dendritic input is required in addition to synaptic input to trigger a plateau potential in segment *i*. For leaf nodes of the dendritic tree we set TD_*i*_ = 0. When both conditions become satisfied, i.e. there is sufficient synaptic and dendritic input, then a plateau potential is initiated. We use 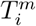 to denote the starting-time of the *m*^th^ plateau potential in segment *i*:

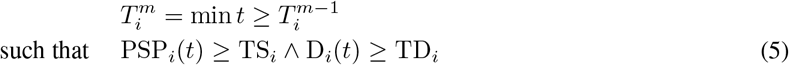

The plateau potential then typically ends at time 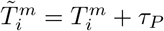 after the fixed duration *τ*_*P*_, unless it is interrupted by shunting inhibition. We formalize this special case as follows: The first inhibitory spike, if any, from neuron *k* ∈ inh_*i*_ at time 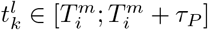 can end the plateau which means 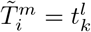 in that case.

We can now define the output of segment *i* as a sequence of binary pulses, the plateau potentials:

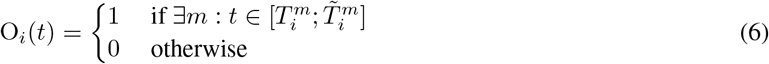

This formalism can be iteratively applied to all segments of a neuron, including the soma that produces a spike event followed by a brief refractory period *τ*_refrac_ instead of a long-lasting plateau potential^4^. A software implementation of the model and all following experiments relies exclusively on these event times as they are the only times at which the state of the neuron can change (see Methods).

Conceptually, each dendrite segment acts first and foremost as a coincidence detector for a volley of synchronized spikes on the fast timescale of EPSPs. On the second, slower timescale of dendritic plateaus each segment is gated by its children in the dendritic tree. In a sentence, the computation of the neuron depends on a sequence of activations of its segments by spike volleys which can be interrupted by shunting inhibition.

## Results

### Dendrites can recognize movement trajectories from place cell activity on multiple timescales

To illustrate how dendritic plateau computation can be used in a close-to-real-world example, consider again the sequential patterns emitted by place cell populations. The location of an animal in its environment is represented by place-cells [1, 2], each of which has a receptive field centered at a specific location. Navigation naturally produces sequential activation patterns as different locations are visited. The timescale of these patterns can be long and is variable because it is directly linked to the movement speed of the animal [3]. This variability in timing is exacerbated during sleep-replay, where the same patterns can be replayed at significantly compressed timescales [63], making a reliable detection particularly difficult. Active dendritic processes likely play an important role in this sequence detection task, since they frequently occur in cortical pyramidal neurons of freely moving rats [64] and have been shown to be selective for specific sequences of synaptic inputs [65]. Plateau computation can allow single neurons to solve this detection problem across varying movement speeds and during replay.

We numerically simulate a rat moving through a small, 2-dimensional environment by generating stochastic paths at varying movement speed (more details in Methods). The environment is tiled by the receptive fields of place cell populations, each 20 neurons strong. These populations emit spike volleys, the magnitude of which increases as the animal gets closer to the center of the respective receptive field (Fig. 2 **(A)**). This encoding could arise from increased firing rates, increased correlation of the spike timings, or both [66]. The task is to recognize whether the animal followed a specific path leading through the receptive fields of three place cell populations in the correct order: from the bottom left (A, in beige) through the center (B, in blue) to the top right (C, in purple). A neuron composed of two sequential dendrite segments and the soma can solve this task reliably if each of the segments receives synaptic input from one of the place cell populations. In this example, a plateau potential is triggered in segment A if TS_*i*_ = 13 or more of the 20 neurons in the associated place cell population fire a synchronous spike volley, which implies that the animal crossed through the corresponding receptive field. This detection is memorized in the elevated membrane potential of segment A for the duration of the plateau. This also raises the resting potential of segment B and enables it to respond to a spike volley from its corresponding place cell population with a plateau of its own. Likewise for the soma, which can finally fire a spike in response to a spike volley from place cell population C. Since each of the successive detections depends on the preceding plateau potential, the neuron only responds if the three stimuli *A* → *B* → *C* occur in the correct order. But because each plateau enables the subsequent segment for a prolonged duration, the exact timing of the next sequence element within this time interval is irrelevant (Fig. 2 (B). Sequence detection via dendritic plateaus thus combines two distinct timescales: the fast estimation of the current location through coincident spike volleys and the slower integration of these events through interacting plateau potentials. The spatiotemporal receptive field of the neuron can thus be very sensitive and responsive to changes in location, but largely invariant to the speed at which the animal travels along the path on a much slower timescale (see supplementary information S1).

**Figure 2:**
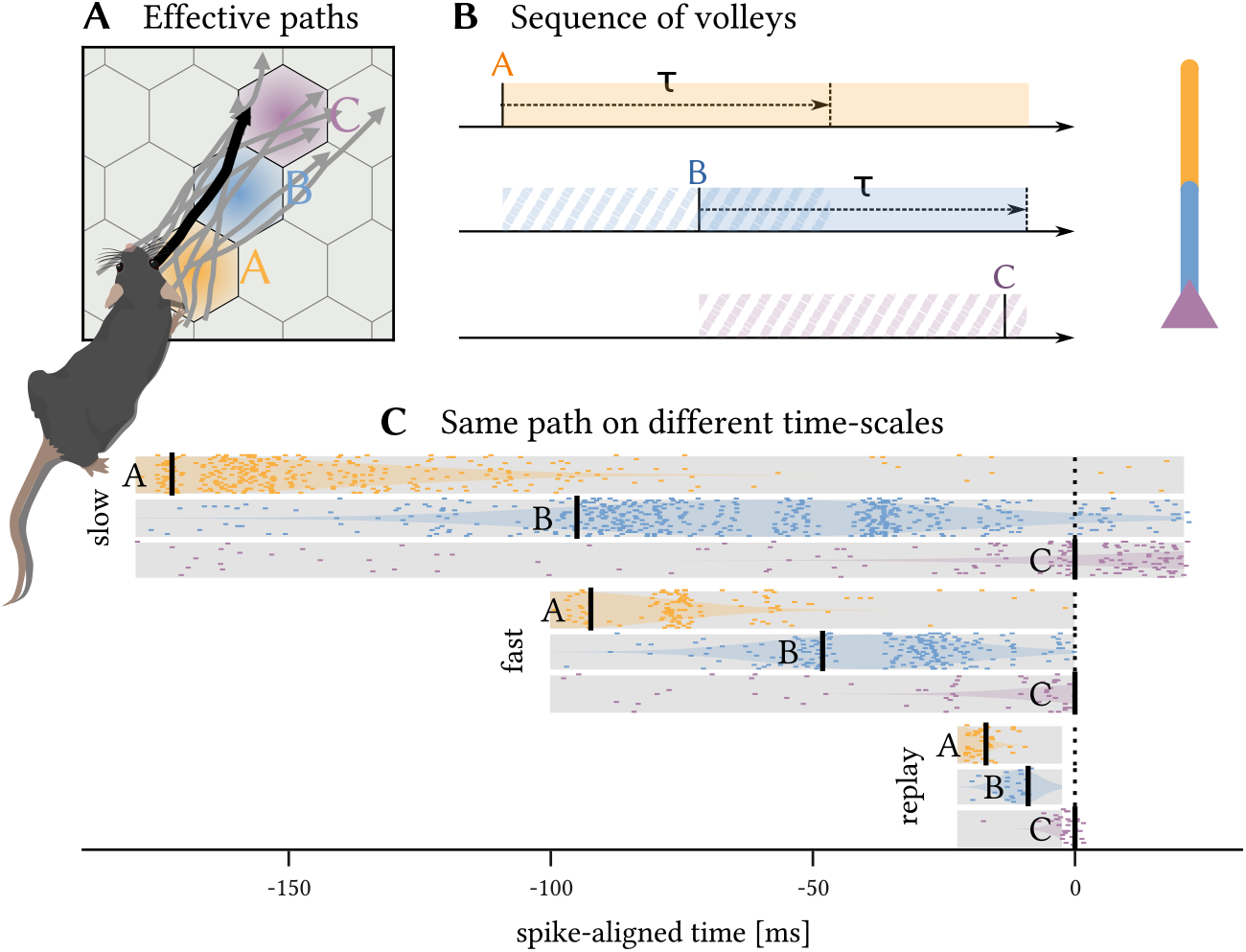
A simple neuron with three dendrite segments as shown to the right of panel **(B)** can detect directed paths on a timescale of 300 ms. **(A)** The receptive fields of place cell populations tile the environment into a hexagonal grid. The populations *A,B* and *C* are each connected to a different segment of the neuron (color coded). Random trajectories are generated through a stochastic process with randomized initial positions, velocities and angular heading to simulate the animal’s movements. A random sample of 10 trajectories that elicit a neural response are shown; all of them follow the same direction through receptive fields *A* → *B* → *C*. **(B)** As the animal passes through a receptive field, the place cell population emits a spike volley that triggers a plateau potential in the connected dendrite segment (shaded regions). This plateau lasts for an extended period *τ*, which provides a broad time window during which a spike volley from the next receptive field can be detected (striped regions). **(C)** The same path (A, solid black line) is repeated at three different speeds (slow movement, fast movement and highly compressed replay). Scatter plots show the spike volleys emitted by the place cell populations; shaded areas indicate the expected magnitude of the volleys over time. For all three timescales, the same neuron detects the sequence *A* → *B* → *C*.

A remarkable property of this mechanism is, that it is also able to decode the same spike patterns during orders-of-magnitude faster compressed replay (Fig. 2 **(C)**). Because compressed re- and preplay mechanisms are implicated in planning and skill learning [67], we expect that this robustness of dendritic sequence processing to compressed pattern representation is critical for theses tasks. However, this timing invariance also implies that techniques to identify spatiotemporal receptive fields via temporal averaging, such as the spike-triggered average [68], are inadequate for neurons with active dendrites. Instead, new techniques are required that identify individual receptive fields of dendrite segments and compare the relative timings of the generated plateau potentials (see supplementary information S2).

### Shunting inhibition can prevent false positives

The mechanism outlined above only relies on excitatory synapses and is able to identify specifically ordered sequential patterns of spike volleys on varying timescales. However, if the same stimuli are frequently repeated in incorrect order, the likelihood of errors increases. To illustrate this, consider the sequence *A* → *B* → *C* and its reverse *C* → *B* → *A*, i.e. the same path but travelled in the opposite direction (Fig. 3). During sleep replay, or while running in a circle, this reverse sequence may be presented multiple times in quick succession. Naturally, the sequence *C* → *B* → **A** → *C* → **B** → *A* → **C** → *B* → *A* also contains the sub-sequence *A* → *B* → *C* (highlighted) which would trigger the neuron. Because the additional excitatory spike volleys cannot prevent the neuron from firing, the repeated presentation of the reverse sequence will therefore lead to a false detection (Fig. 3 **(A)**). Anti-patterns such as this are an inevitable side-effect of the desirable timing-invariance of sequence detection.

**Figure 3:**
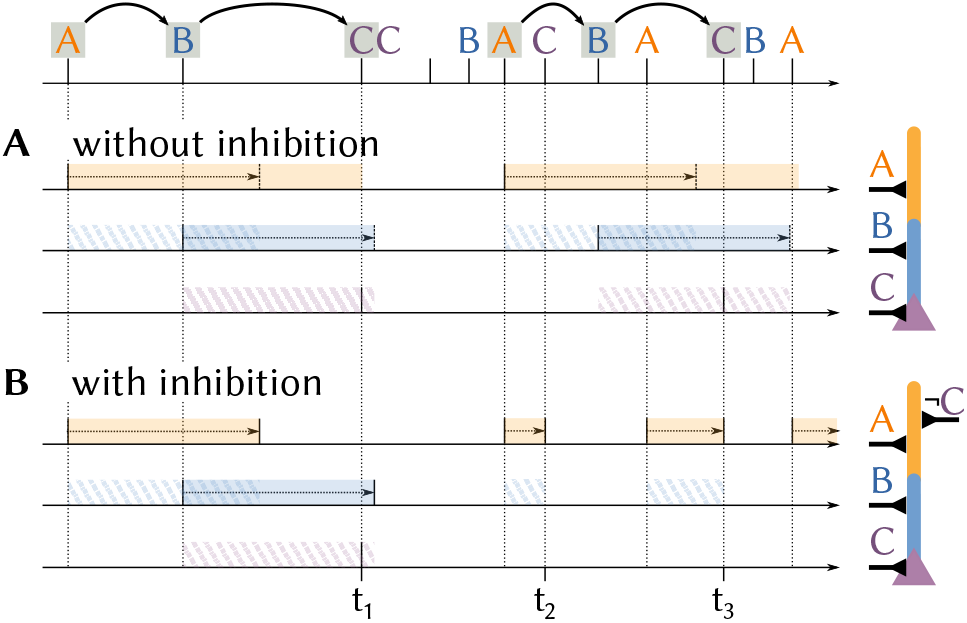
Shunting inhibition can prevent false detections. A neuron receives a sequence of spike volleys from three populations *A, B* and *C*. **(A)** A neuron with a chain of dendrite segments A and B and soma C fires whenever they are activated in the correct order *A* → *B* →*C*, for example at time *t*_1_. This also results in a false detection at *t*_3_ if the desired sequence *A* → *B* → *C* is contained in fast repetitions of the undesired sequence *C* → *B* → *A*. **(B)** By adding shunting inhibition, input for segment *C* also stops the plateau at the first segment *A* at *t*_2_ and *t*_3_ which prevents the false detection.

Shunting inhibition can prevent such false positives and restore the neuron’s high selectivity to sequence order in these situations (Fig. 3 **(B)**). Consider the same situation as before, but now additional inhibitory synapses from population *C* can disrupt plateaus in the neuron’s first segment. If a spike volley from *C* immediately follows a volley from *A*, the previously generated plateau in the first segment is stopped and sequence detection has to start anew with a novel detection of a spike volley from population *A*. However, if populations *A, B* and *C* all fire in order the neuron will fire a spike before the inhibition of the first segment takes effect. Inhibition therefore acts as an important complementary mechanism for dendritic plateau computation and can “veto” anti-patterns from erroneously activating sequence detecting neurons.

Unlike excitatory spikes, which need to be synchronized into spike volleys to efficiently drive plateau generation, shunting inhibition can disrupt a plateau potential at any point.

### Probabilistic and graded plateau responses increase information content

In the previous example, each place cell population codes for a single location. The magnitude of the spike volley encodes the proximity to the receptive field center, or, in other words, the evidence for the fact that the animal was close to the specified location. If the spike volley is transmitted deterministically (*p* = 1.0), the receiving dendrite segment detects it if and only if it exceeds a hard threshold at which a plateau is generated (Fig. 4 **(A)**). If the spike volley is instead transmitted stochastically (*p <* 1.0), the probability of generating a plateau is directly proportional to the spike volley magnitude (see Methods). The response of an individual dendrite segment to one spike volley is of course binary in both cases, but if we assume that many segments respond to the same spike volley with stochastic synapses, then the total number of emitted plateau potentials becomes a smooth, sigmoidal, graded function of the volley’s magnitude (Fig. 4 **(B)**). In an ensemble of neurons with dendrite segments that respond to the same input probabilistically and independently due to unreliable synaptic transmission, the number of active plateau potentials can therefore encode the magnitude of the received spike volleys.

**Figure 4:**
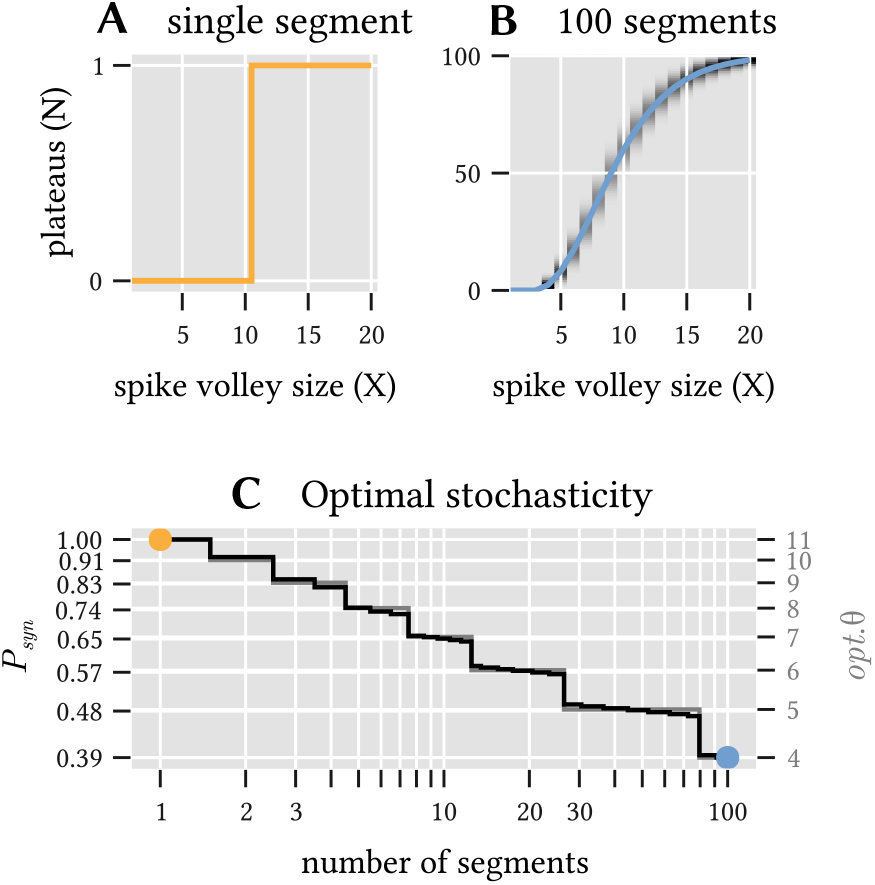
Unreliable synapses can improve information transmission for ensembles of dendrite segments. **(A)** A single dendrite segment with 20 inputs and deterministic synapses (transmission probability *P*_syn_ = 1.0 and threshold *θ* = 11) provides a binary classification (1 bit) of the magnitude *X* of the incoming spike volley. **(B)** The expected number *N* of plateaus (solid line) in an ensemble of 100 segments with stochastic synapses (*P*_syn_ = 0.39 and *θ* = 4) is a continuous function of the volley magnitude *X* and thus conveys more than 1 bit of information. Grey shading indicates the probability distribution around this mean value. **(C)** Combinations of *θ* and *P*_syn_ that maximize the mutual information between the magnitude of the incoming spike volleys and the number of generated plateau potentials for a varying number of independent segments. Note that increased stochasticity is beneficial as the ensemble size increases.

How well this encoding works can be quantified by the mutual information between spike volley magnitude and number of triggered plateaus. While there is clearly no benefit to synaptic stochasticity if only a single segment receives the spike volley, multiple dendrite segments can in fact benefit significantly from stochastic synaptic transmission. For example, a synaptic transmission probability as low as *p* = 0.39 maximizes information transmission for an ensemble of 100 dendrite segments that receive spike volleys from a population of 20 neurons. In this case, unreliable synapses allow an ensemble of otherwise identical deterministic neurons to exceed the information capacity of any individual neuron (Fig. 4 **(C)** and Methods).

In our model, consecutive segments gate consecutive plateaus. In the path-detection example above, three consecutive spike volleys in a sequence must all be detected for the neuron to fire a spike. The probability of the spike response is therefore the product of all three plateau probabilities and thus directly proportional to the combined evidence for all sequence elements as encoded by the corresponding spike volleys. This affords another interpretation of the neuron’s behavior: the probability to fire a spike encodes how closely the animal passed each of the receptive field centers along the path. An ensemble of neurons is therefore be able to report not only if the animal followed some desired path, but also how closely its chosen path came to the three receptive field centers.

In the context of sequence detection, the stochasticity of synaptic spike transmission serves one other important purpose. Since plateau potentials endow the neuron with a long lasting internal state, any neuron that is already engaged in the detection of a sequence is not ready to detect another until its plateau potentials have subsided. This implies, however, that a relevant sequence might not be detected at all if it is directly preceded by stimuli that partially activate the neuron. This also imposes a strict limit on the maximum rate at which any sequence could be detected: the inverse of the plateau duration, which is independent of the actual length of the sequence. If synapses would transmit spikes deterministically, even an ensemble of neurons would suffer from the same problems. Neurons, depending on initial condition and synaptic connections, will operate in lockstep and detect the same instances and partial sequences of the target sequence. Hence, they may also fail to detect the target sequence if neurons in the ensemble are partially engaged in plateaus. Synaptic stochasticity helps by temporally decorrelating the individual detectors such that some subset of neurons is always available.

### Stochastic spike volleys can encode events in multi-dimensional feature space

So far, we only considered sequences of discrete events, such as an animal traversing specific receptive fields in order. In many situations, however, the individual sequence elements may not be as simple and binary as that. Instead, each sequence element may itself be encoded by a combination of more general features. We find that the stochastic plateau generation also provides a simple way to represent such multi-dimensional feature combinations, which allows the detection of yet more complex patterns.

We identify each input feature with a spike-volley from a specific randomly selected subset of a much larger neuron population. These subsets can overlap for different features, but the expected overlap is small for sufficiently sparse subsets. By stochastically co-activating different fractions *p*_*i*_ of neurons from each of these sub-populations *i*, a single spike volley can thus encode various combinations of features. This approach is closely related to hyper-dimensional computing [69]; see Methods for a formal derivation.

Suppose we are interested in combinations (*p*_1_, *p*_2_, *p*_3_) of three features *P*_1_, *P*_2_ and *P*_3_, where for simplicity 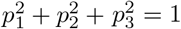 is normed. The possible feature combinations thus span a continuous two-dimensional manifold, a sector of the surface of the unit sphere, embedded into the high-dimensional space of the neuron population’s spiking output. Figure 5 **(A)** shows an approximately distance-preserving projection of this manifold.

**Figure 5:**
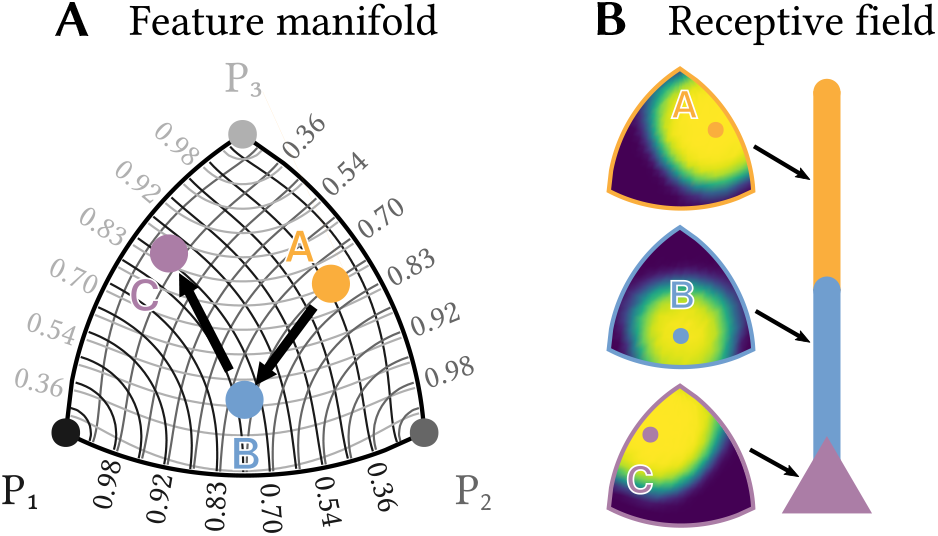
Individual neurons can detect sequences of feature combinations. Consider a sequence of the three elements *A, B* and *C*, each of which is a different normed combination of the three basis features *P*_1_,*P*_2_ and *P*_3_ and therefore lies in a two-dimensional manifold. **(A)** shows an approximately distance preserving projection of that manifold. Contour lines show the similarity of each point to the three basis features. **(B)** Each of a neuron’s segments can be tuned to respond only to a specific combination of basis features by a corresponding choice of synaptic transmission probabilities. Heatmaps show the resulting receptive fields for a neuron designed to detect the sequence *A* → *B* → *C*.

Within this manifold, we want to detect a specific sequence *A* → *B* → *C* of three different combinations of the basic features. The same kind of neuron as described above can solve this task, if the transmission probability of each synapse is tuned to the desired strength *p*_*i*_ of the corresponding features. Each dendrite segment can then selectively respond with a plateau potential to only one specific combination of features (see figure 5 **(B)**), which allows the neuron to detect the sequence *A* → *B* → *C*. Instead of changing the magnitude of EPSPs for different synapses, this approach works by tuning synaptic transmission probabilities.

### Populations of plateau computing neurons improve sequence detection

The sequence detection capabilities of individual neurons can be extended to networks. As a sequence of interest gets longer and comprises more elements, it seems increasingly unlikely that the brain would rely on a single highly complex and specific neuron, or an ensemble of such neurons, to detect it. It is much more likely that multiple neurons code for different, shorter sequences – features – that occur in different combinations as parts of longer sequences. In analogy to the original Perceptron model [70], these features can be task-independent and chosen at random. To detect a specific long sequence of interest, a neuron would then only need to detect the right combination of a few of these features in correct order. A concrete example is the ten element sequence *D* → *B* → *C* → *A* → *F* → *E* → *D* → *H* → *B* → *F*. It contains an enormous number of possible features such as (*D* → *B* → *C*), (*C* → *A* → *E*), (*D* → *C* → *E*) and so on, each of which might be detected by one random sequence detection neuron with just two dendrite segments and soma (see Fig. 6**(F)**-**(H)**). If many of these short features are present in the stimulus in the correct order, which can be recognized by a specialized sequence detection neuron, this provides strong evidence for the long sequence.

**Figure 6:**
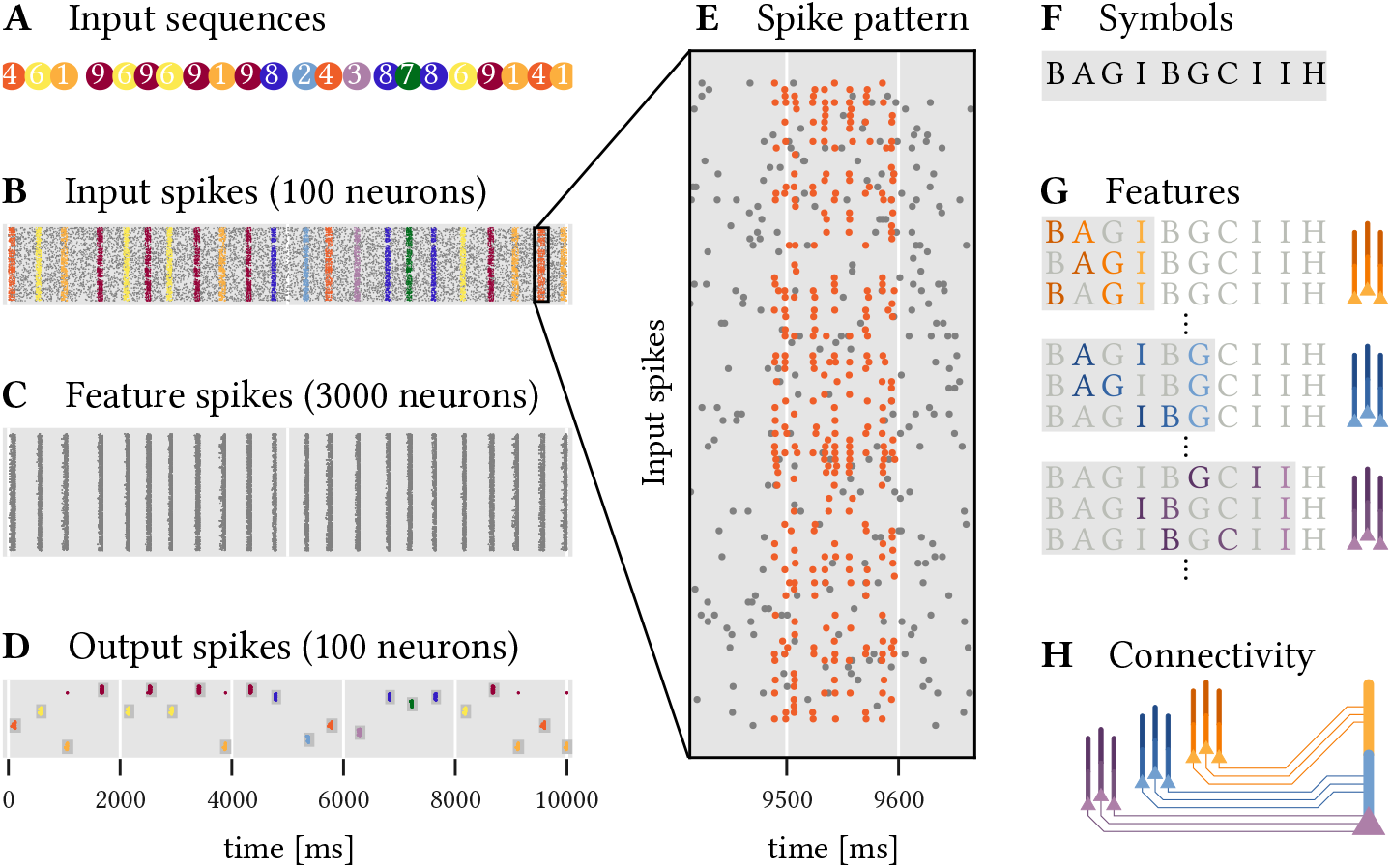
A two layer network of sequence detecting neurons with dendritic plateaus recognizes complex, overlapping, and long sequential patterns. 10 random sequences (1 through 10) of 10 random code-words (*A* through *J*) have to be detected. **(A)** The different target sequences are presented in random order (color coded). **(B)** Each of these instances results in a temporal pattern of spikes across 100 input neurons. Noise is modeled by additional spikes (gray dots). See also **(E)** for a close-up of sequence 5, which is composed of the symbols shown in **(F). (C)** 3, 000 neurons in a first layer each detect a random sequence of three symbols from the input spikes. **(D)** 100 Neurons in the output layer detect specific sequences of spike volleys from the hidden layer, i.e. sequences of sequences of symbols. This leads to a rather reliable detection of the ten different target sequences by the corresponding output neurons. **(G)** Sub-populations of neurons in the hidden layer that code for different sequences ending in the same symbol fire synchronized spike volleys. **(H)** Neurons in the output layer can detect specific sequences of these spike volleys that are characteristic of the respective target sequence.

We set up a simple experiment to demonstrate how effective this strategy can be. The task is to classify ten different sequences of spike volleys (labeled 1 to 10, see Fig. 6**(A)**) that are generated by a population of 100 input neurons. Each of these sequences is a series of ten randomly selected symbols with randomized timing. There are ten symbols (labeled *A* to *J*) in total and each is represented by a spike volley from a randomly selected subset of 30 input neurons. The same symbols may appear multiple times within a sequence and are shared across the different sequences which makes classification challenging. In addition to these stimuli, the input neurons also generate spikes at random to emulate a backdrop of noise. Fig. 6**(B)** shows the resulting spikes from the input population, and Fig. 6**(E)** zooms in on a single example of such a sequence.

To solve this task, we construct a network of simple sequence detection neurons arranged into two layers. All of the neurons comprise just two dendrite segments and the soma in series, just like in the examples above, and are therefore able to detect sequences of up to three elements. In the first hidden layer of the network, 3000 neurons each detect a random sequence, or feature, composed of three symbols (Fig. 6 **(C)**). These features are drawn entirely at random from the set of all possible symbols^5^. Each segment and the soma receives input from the 30 input neurons that represent the associated symbol. A second layer of 100 output neurons (ten for each target sequence, see Fig. 6 **(D)**) uses the spikes coming from the hidden layer to classify the ten different target sequences. To achieve this, each output neuron needs to detect a sequence of feature-combinations that is characteristic for the target sequence of interest. Each segment and soma of an output neuron thus detects a combination of relevant features, all of which terminate in the same symbol and thus produce a synchronized spike volley once that symbol appears in the target sequence. Here, we use a simple algorithm to select these feature combinations (see Methods), but the same might be realized through a combination of structural and homeostatic plasticity in the brain. Now, whenever the target sequence is presented, the corresponding feature detectors in the hidden layer fire in order, which successively activates the dendrite segments of the output neurons, ultimately leading to spikes that signals a detection. Despite noisy inputs and timing variability in the spike patterns, the results are remarkably robust and each target sequence is reliably detected. Because the feature detectors in the hidden layer are agnostic to the target sequences, this scheme can be extended to another target sequence by adding an additional output neuron to detect the right sequence of feature combinations.

## Discussion

Mounting biological evidence suggests that dendritic function is critical to neural computation. We therefore constructed a new model of dendritic computation that relies primarily on the generation and interaction of dendritic plateau potentials. It explains conceptually how individual neurons with segmented dendritic trees can robustly detect long sequences of spike volleys in a particular order – even if they are presented across vastly different timescales. We also showed that stochastic synapses can turn the all-or-none response of plateau potentials into a more informative graded response. This allows individual neurons to detect complex sequences of stimuli that are themselves combinations of multiple features. Stacking such sequence detection neurons as features into a larger population further expands their ability to detect very long sequential patterns; even when based on generic, randomly generated feature detectors.

Our work is closely related to the hierarchical temporal memory model of neural computation [71] in which active coincidence detection in dendrites generates neural UP states [23]. In that model, sequence detection is realized on the network level through the interaction of neurons in UP states. Other related work also makes use of a single, active dendrite compartment to facilitate learning of temporal patterns via synaptic plasticity [72]. In contrast, our model explicitly includes both the fast timescale of passive membrane potential dynamics and the much longer timescale of plateau potentials. This allows an individual neuron to cope with significant timing variability in sequential patterns. Further, the structure of the neural dendrite, its functional compartmentalization, and the location of synapses determine the neuron’s ability to detect specific sequences in our model. This provides a different perspective on the functional relevance of the observed diversity of dendrite morphologies and their potential ability to adapt compartmentalization in cortex [50].

We limited our discussion here to dendrites composed of a single chain of segments, but future work should also consider more intricate branching patterns that can enable individual neurons to detect more complex sequential patterns such as “A or B and then C” (see supplementary information S3). Since plateau generation requires highly correlated synaptic input, we also rely on spike volleys as the primary unit of information transmission. Although new statistical techniques are being developed to detect such volleys in-vivo, there are competing ideas about the mechanisms that could cause this synchronization. A particularly interesting prospect is the emergence of stimulus-dependent synchronization from recurrent activity [73]. Our hypothesis that plateau potentials are an integral part of neural computation is supported by a growing number of recent studies that show long-lasting calcium signals in dendrites to be associated with task relevant information in a variety of experiments [21, 22, 74].

Here, we focused entirely on the role of dendritic plateaus for computation. Future work should also address their implications for learning. For example, it is still unclear how dendrites can learn to detect specific sequences based on local plasticity mechanisms alone. This is especially important, because localized plateau potentials and the resulting long-lasting high Ca^2+^ concentrations appear to be the primary drivers of synaptic plasticity [75]. This is at odds with most current learning algorithms for spiking neurons, which instead rely on short somatic feedback signal such as backpropagating action potential [76].

Other forms of plasticity are even more relevant in the context of our proposed model. For example, since the location of a synapse on the dendrite matters, structural plasticitity [77] is decisive for the neuron’s function. The localized coincidence detection requires homeostatic processes to adjust synaptic transmission probabilities [78, 79] in order to appropriately balance the excitability threshold and synaptic input. And lastly, a recently proposed mechanisms for tuning transmission delays through controlled (de-)myelinization [80] might allow neuron populations to synchronize their spike volleys more precisely.

Beyond its role as a candidate mechanism for sequence processing in the brain, dendritic plateau computation may also have applications outside of neuroscience. In particular, the simplicity of our proposed mechanism lends itself to a hardware implementation that uses asynchronous communication and complex dendrite structures, rather than larger networks, to boost computational efficiency. The prospect of energy efficient dendritic computation has also motivated others to research potential implementations in neuromorphic hardware. For example, Intel’s Loihi chip [81] and the DYNAPSE architecture [82] already support some forms of active non-linear processing in functionally isolated dendrite segments. Our model provides a new perspective on how these existing capabilities could be utilized for computation and extended in future neuromorphic technologies.

## Materials and Methods

### Simulation framework for dendritic plateau computation

All simulations are implemented in a custom package developed in the Julia programming language [83], publicly available via the code repository hosted at https://github.com/jleugeri/DPC.jl. The simulator implements the neuron model outlined in this paper using a fast and extensible event-based formalism. All experiments and configuration files can be found in the examples subfolder of the repository. Further documentation of the simulator, its interfaces, and implementation details can be found there as well.

### Implementation of the navigation experiment

To simulate the stochastic movements of an animal in a two-dimensional environment, random paths are generated with time-varying location *l*(*t*) = (*X*(*t*), *Y* (*t*)) ∈ ℝ^2^ as solutions of the following system of stochastic differential equations:

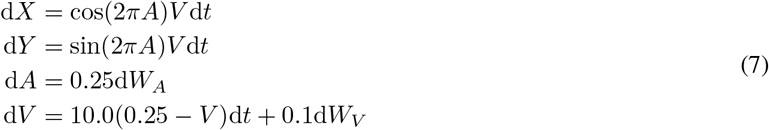

*A* represents the angular heading of the animal, *V* represents its velocity in m s^−1^ and *W*_*A*_, *W*_*V*_ represent independent standard Brownian motion processes. Each path is generated with a randomized initial position within a rectangular domain of 10 cm × 9.5 cm, a random angular heading and a random velocity according to the marginal stationary distribution of *V* in the equation above, and is simulated for a fixed duration of 200 ms. Three populations of place cells, each 20 neurons strong, are centered on a hexagonal grid with center-to-center distance of *r* ≈ 2.9 cm. Each population randomly emits spike volleys following a homogeneous Poisson process with rate *λ* = 250 Hz. The magnitude of each spike volley is determined by the population’s mean activity at the time which depends on the animal’s location within the environment through a receptive field tuning curve. The tuning curves model the probability of each individual neuron within the population to participate in a given spike volley by the bell-curves 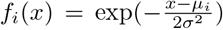 with coefficient *σ* = 9.7 mm, centered on the tiles of the hexagonal grid. The total number of spikes emitted during a volley from population *i* at time *t* is therefore a random variable distributed according to a Binomial distribution with population size *n* = 20 and probability *p* = *f*_*i*_(*l*(*t*)). Additionally, each neuron in the population emits random spikes at a rate of 10 Hz to emulate background activity.

Each of the simulated neuron’s dendrite segments receives spiking input from the 20 neurons of one place cell population and requires 13 coincident spikes to trigger a plateau potential. The three segments are connected in a chain that requires sequential activation by spike volleys from the input populations in correct order to fire a spike. A random path is considered to be accepted by the neuron if the neuron responds with a spike at any point in time during the corresponding simulation run.

### Implementation of the stochastic plateau generation experiment

For a single dendrite segment with *K* = 20 stochastic synapse, each with transmission probability *P*_syn_, the total number of transmitted spikes for an incoming volley of *X* spikes is a conditional random variable *S*|*X* ∼ Binomial(*X, P*_syn_). Whether the segment fires a plateau in response to the volley (*Z* = 1) or not (*Z* = 0) is another conditional random variable *Z*|*X* ∼ Bernoulli(*P*_*Z* | *X*_), where *P*_*Z* | *X*_ = *P* (*S* ≥ *θ*) = 1 − *F*_*S* | *X*_ (*θ* − 1) is the probability to exceed a fixed threshold *θ*. In an ensemble of *M* i.i.d. segments that receive the same spike volley as input, the number of triggered plateaus is then again a conditional random variable *N*|*X* ∼ Binomial(*M, P*_Z|X_). If the magnitude of the spike volleys *X* ∼ *P*_*X*_ is chosen at random from some distribution *P*_*X*_ (here a discrete uniform distribution on [1, 20]), the amount of information conveyed by the number of triggered plateaus can be computed as

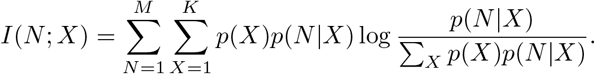

For different numbers of segments *M* from 1 to 100, we perform a grid search over the parameters *P*_syn_ and *θ* and identify the optimal parameter combination that maximizes the mutual information. The two parameters are almost perfectly correlated (see Fig. 4C), with an optimal synaptic transmission probability of 1 (i.e. deterministic synapses) and a threshold of 11 for *M* = 1, and a transmission probability of only 0.39 (i.e. highly stochastic synapses) with a correspondingly lowered threshold of 4 for *M* = 100 segments. For these two extreme cases (see Fig. 4A and B), we vary *X* from 0 to 20 and plot the expected number of plateaus (solid lines) as well as the conditional probability *P* (*N* |*X*) (gray heatmap).

### Implementation of the multi-dimensional feature space experiment

For a neuron population of *N* neurons (here *N* = 1000), we can collect the indices of the neurons associated with a particular feature *i* in a sparsely populated binary vector *v*_*i*_ ∈ {0, 1}^*N*^. The degree to which feature *i* matches the current stimulus *j* can be encoded by the probability 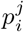 that an input neuron coding for feature *i* would actually participate in a spike volley for *j*. Since the same input neuron *k* can be associated with multiple (here *M* = 3) features *i* ∈ {1, 2, 3}, its total probability to fire a spike for stimulus *j* with coefficients 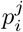 is

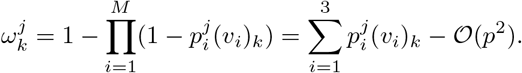

For small *p*_*i*_, the expected input vector *ω*^*j*^ thus approximates a weighted linear combination of the basis vectors *v*_*i*_ with coefficients 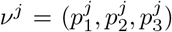. We require that 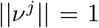. Using the same equation, we can fix a set of target coefficients 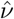 with 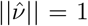 and compute the optimal weight vector 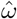. The total number *X* of spikes received for stimulus *j* by a endrite segment with synaptic transmission probabilities 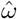 is then a random variable with expected value 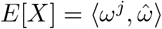. The expected value is maximized for 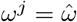, i.e. when input and weight vector are perfectly aligned. Since both *ω*^*j*^ and 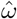 are normalized, they lie on a two-dimensional manifold (the positive sector of the surface of a unit sphere). The probability that *X* exceeds a segment’s threshold *θ* for a stimulus *j* and thus causes a plateau is a radially symmetric function on this manifold. Fig. 5B shows an approximately distance preserving projection of this probability distribution for three different weight-vectors, the coefficients of which correspond to the coefficients *ν*^*A*^, *ν*^*B*^ and *ν*^*C*^ of the three sequence elements *A, B* and *C*, respectively. See supplementary material for details on the used projection method.

### Implementation of the network experiment

There are 3000 neurons in the hidden layer, each of which codes for a different random sub-sequence (three elements long) of any of the ten target sequences (ten elements long). The synaptic transmission probability is 0.5, and 8 coincident spikes are needed to trigger a plateau (or somatic spike) in the hidden layer. Of the 100 output neurons, each segment (and the soma) receives synaptic input from a different subset of the hidden layer’s neurons. Since our model relies on synchronized spike volleys, these inputs are chosen according to a simple heuristic algorithm: Each segment *i* (and the soma) of an output neuron is associated with a specific index *X*_*i*_ of the neuron’s target sequence in appropriate order, i.e. 1 ≤ *X*_*i*_ *< X*_*j*_ ≤ 10 for *i < j*. Now consider only the slice of the neuron’s target sequence that includes the first *X*_*i*_ elements. Any neuron in the hidden layer is connected to segment (or soma) *i*, if it codes for a feature that occurs as a sub-sequence of this slice and ends with the symbol at index *X*_*i*_. All neurons connected to the same segment thus code for features ending in the same final symbol. Therefore, they form a population that fires a spike volley when triggered by this final symbol. The magnitude of the resulting volley depends on the number of features that were detected by the population for the current stimulus. Since the size of each of these input populations is random, the synaptic threshold for plateau or spike generation in a segment or soma is dynamically set to 40% of the number of synapses.

## Footnotes

2 We avoid the term “compartment” to prevent confusion with the concept of multi-compartment neuron models, which are commonly used as a spatially discretized solution to partial differential equation models of neurons.

3 We assume that spike arrival times 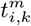 are at least *τ*_*E*_ apart.

4 In addition to the forward-propagation of membrane potentials that we focused on so far (i.e. from child branches to the parent), the reverse direction typically has an even stronger effect — strong enough for the parent segment to depolarize its child segments by itself. To capture this effect, we recursively define that a neuron segment *k*’s membrane potential *V*_*k*_(*t*) = *O*_*k*_(*t*) ∨ *V*_*i*_(*t*), *k* ∈ child_*i*_ is depolarized whenever either the segment itself or any of its ancestors produces a plateau potential. However, we focus on the forward-propagation of plateau potentials and their role for dendritic computation here.

5 We discard sequences that are not a sub-sequence of any target sequence. We call a sequence *X* a sub-sequence of another sequence *T* if all elements from *X* also occur in *T* in the same order.

