## Supplementary Information for "Dendritic plateau potentials can process spike sequences across multiple time-scales"

June 21, 2022

### An approximately distance preserving projection algorithm (used in Fig. 5)

In the experiment shown in Fig. 5 of the main text, we work with coefficient vectors $\nu_i \in \mathbb{R}^3_+, ||\nu_i||_2 = 1$. These vectors all live on a two-dimensional manifold, specifically on the intersection of the surface of a unit sphere with the cone $\mathbb{R}^3_+$. In our example, we are interested only in those vectors that form a normed positive linear combination of three basis vectors inside this manifold, where these basis vectors are not necessarily orthogonal to each other. On the surface of the unit sphere, these basis vectors would form a triangle, with any of the combinations falling inside. Since we make an argument about detecting points on the manifold based on their "similarity" to other points on the manifold, which we measure by the geodesic / great circle distance, we would like to visualize this manifold in a way that preserves these distances. Therefore, we would need a distance preserving (metric) projection of a part of the sphere's surface onto 2D space. Unfortunately, as millennia of cartography can attest, this is impossible; instead, we have to use an approximation that relaxes some constraints. The most well-known projections either preserve angles (conformal maps), area (equal-area maps) or distance to one reference point (azimuthal equidistant), two reference points (two-point equidistant maps) or the meridian [1]. However, neither of these projections is ideal for our application, since we would like to preserve as well as possible the distances to the *three* corner points of the triangle in a symmetric way. The only projection that aims to achieve this, to our knowledge, is a heuristic method called *Chamberlin's trimetric projection* [2] that was developed for and used by the National Geographic society. Since this method is heuristic and no standard implementation is available, we use a custom projection algorithm instead, which we summarize here.

First, we fix the 2D projections $\hat{P}_1$, $\hat{P}_2$ and $\hat{P}_3$ of the three corner points $P_1$, $P_2$ and $P_3$, such that they form a triangle, where the lengths of the sides are proportional to the geodesic distances between the points on the unit sphere. Then, for a point $X$ that is to be projected, we determine the geodesic distances

$d_1, d_2, d_3$ between $X$ and $P_1$, $P_2$ and $P_3$. We now wish to find a point $\hat{X}$ in 2D, where the relative distances to the corner points are preserved, i.e. $||\hat{X} - P_1||^2 : ||\hat{X} - P_2||^2 : ||\hat{X} - P_3||^2 = d_1 : d_2 : d_3$. This gives us two constraints $||\hat{X} - P_1||^2 d_3^2 = ||\hat{X} - P_3||^2 d_1^2$ and $||\hat{X} - P_2||^2 d_3^2 = ||\hat{X} - P_3||^2 d_2^2$, each of which constrains the set of possible solutions to either a straight line or a circle. If these constraint sets do not intersect, then the projection has no solution. This is the case only for some regions of the sphere's surface outside the triangle spanned by the corner points, and is therefore of no concern here. If the constraint sets do intersect, there may be one or two solutions. If there are two solutions, the solution with the smaller sum of distances to the projected corner points is chosen (the other solution then corresponds to the projection of a point on the opposite side of the sphere). For a full derivation and implementation, we refer to the code repository hosted at https://github.com/jleugeri/DPC.jl and the documentation and comments within.

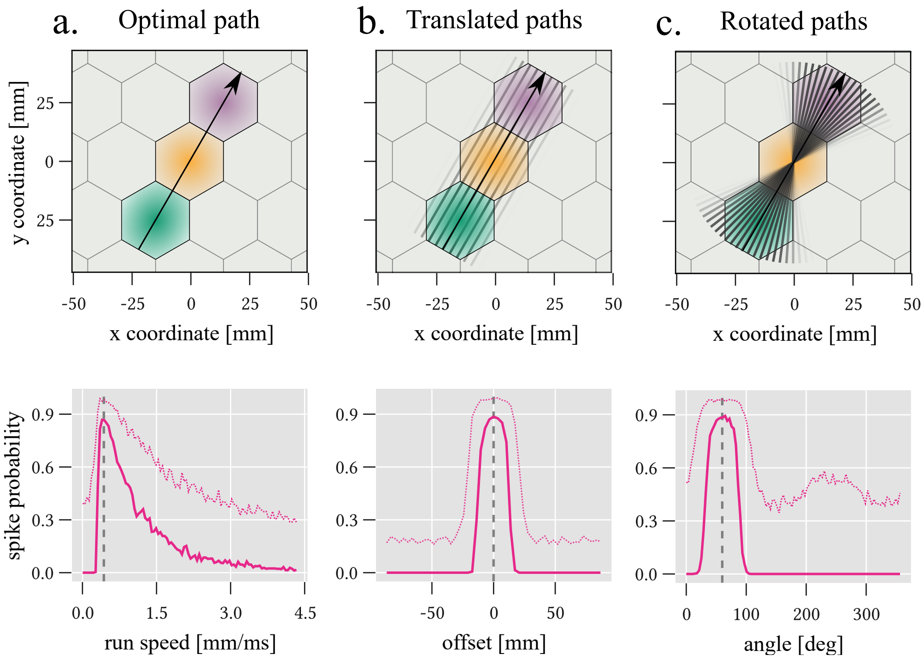


Figure 1: Specificity and invariance of dendritic sequence detection. We systematically varied the optimal path through three place cell populations to characterize the probabilistic response of a neuron that uses dendritic sequence detection over 500 trials. All transmission probabilities are 0.5. Panel a. shows this response when the movement speed along the trajectory is varied. As soon as a minimal run speed is achieved the neuron reliably recognizes the sequence of locations. Depending on the spike volley threshold to generate a plateau ($\theta = 6$ solid line, $\theta = 3$ dashed line), this probability falls off at different rates as the animal moves faster. As seen in the main text, this is not because of inherent speed sensitivity of the plateau mechanism, but because slower paths generate multiple volleys that can be detected, increasing the plateau probability. Panel b. shows the much higher sensitivity of the mechanism to spatial shifts out of the receptive field, both when the threshold is high or low (speed $v \approx 43\,\mathrm{cm\,s^{-1}}$). The spatial selectivity to rotation of the optimal path (Panel c.) is also highly sensitive in the higher threshold case. If the threshold is lowered, this specificity starts to blur out, because the path is guaranteed to go through the center of the orange receptive field.

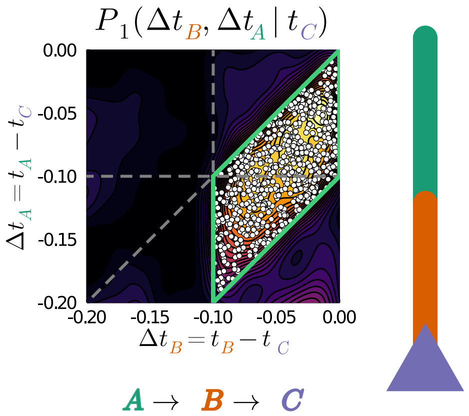


Figure 2: Relative plateau times as an indicator for sequential computation. We drove a sequence detecting neuron with three segments (threshold $\theta = 5$) each receiving input from a population with 20 neurons and constant Poisson rate of $r = 25$Hz for the duration of 1h. For each somatic spike at $C$, we record all plateau times at $A$ and $B$ in the previous 200ms and estimate the joint distribution of over relative plateau times $\Delta t_A$ and $\Delta t_B$ with respect to how much before the somatic spike each occured (heatmap). If we recorded one plateau event for each segment, we called these pairs unambiguous and plotted each event as a white dot. Because the fundamental mechanism of plateau computation in a dendritic tree such as this one is the sequentiality of these events within one plateau time of each other (was 100ms in this experiment), all these unambiguous pairs fall into a parallelogram indicative of exactly this timing relationship. With plateau duration and event time recorded in experiment, a similar method could be used establish the sequential nature of plateau computation in real neurons.

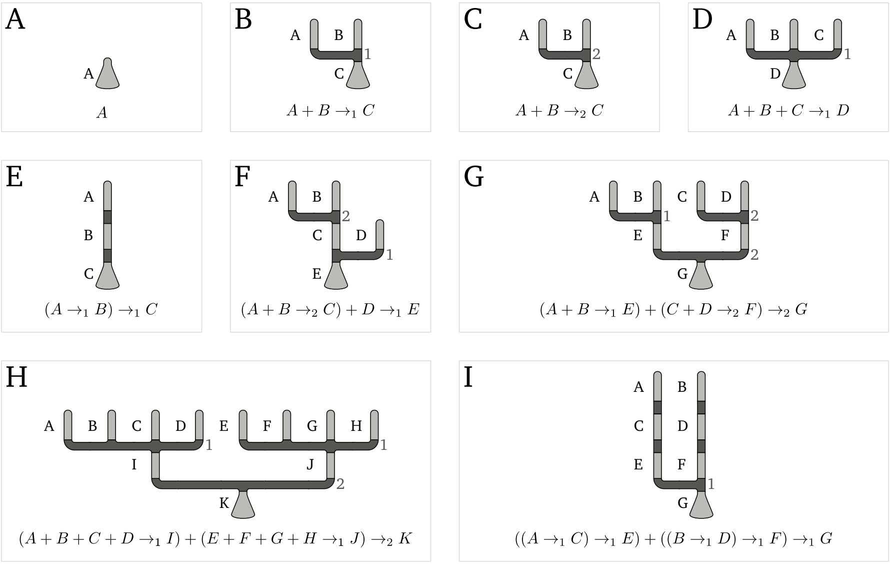


Figure 3: Logical computation on sequence elements in dendritic trees. Consecutive gating of plateau events in compartmentalized dendritic trees may also extend to more intricate branching structures and allow for more complex functions to be computed in active dendrites. Here, the additional mechanism used is the dendritic threshold. This can lead to one or more child segments of any specific dendrite segment required to be active before the parent segment can emit a plateau. To illustrate which kinds of functions may be computed, we use the subscripted $\to_i$ arrow to signify the dendritic threshold. Without considering different weights for different dendrite segments, we can shorten expressions such as "plateau A + plateau B ¿ 1, and then plateau C" (Panel B) to simply say "A or B, then C". With a threshold of 2, this description of the neurons detected pattern becomes "A and B, then C'. The remaining panels give an indivation of how intricate these expressions can become.
